# Multi-region biopsies and patient-derived neurosphere cultures reveal spatial divergence in glioblastoma

**DOI:** 10.64898/2026.04.07.717044

**Authors:** Roberto Salatino, Jacob Geisberg, Arantxa Romero-Toledo, Benjamin Oakes, Jerome C. Nwachukwu, Dobeen Hwang, Cristina Vincentelli, Oszkar Szentirmai, Thomas O. McDonald, Kendall W. Nettles, Franziska Michor, Michalina Janiszewska

## Abstract

Intratumor heterogeneity (ITH) is one of the main reasons for the lack of effective targeted therapies for glioblastoma (GBM). Imaging-guided surgical navigation allows for tumor-wide sampling to account for variation across distant regions of the tumor, but typical drug screening is performed on cell lines derived from a single biopsy and does not account for GBM heterogeneity. Here we profiled matching MRI-guided multi-region primary tumor biopsies from 6 GBM cases (n=40 biopsies) and corresponding neurosphere cultures (n=30) derived from these spatially distinct tumor samples. We found that *in vitro* cultures derived from distinct regions of the same tumor display divergent phenotypes, proliferative capacity and ability to accumulate 5-aminolevulinic acid, used to visualize cancer cells during surgery. The differential drug response of the multi-region neurospheres remains linked to the gene expression of the original tumor biopsies. Thus, studies with multiregion-derived neurospheres are essential to faithfully model GBM ITH for therapeutic testing.

**KEY POINTS:**

- Multi-region biopsy-derived neurospheres represent distinct spatial locations in the GBM tumor.
- Cultures derived from different regions of the tumor retain phenotypic diversity.
- Parental biopsy phenotype predicts drug response better than to in vitro phenotype.

**IMPORTANCE OF THE STUDY:** Cell lines developed from spatially distinct regions of glioblastoma capture its intratumor heterogeneity. We show that while the transcriptional output of these cell lines is not connected to their spatial origin, their drug response can be linked to it. Thus, spatial heterogeneity reflected in our neurosphere collection provides a new paradigm for drug screening in these highly heterogenous and difficult to treat tumors.

## INTRODUCTION

Advances in tumor profiling have enabled the development of precision cancer medicine and have drastically improved survival for many cancer patients^1^. Identification of tumor subtypes based on genomic and transcriptomic profiling has emerged as a stratification tool for clinical trials for specific molecular markers across tumor types. In the case of glioblastoma (GBM), the most aggressive brain tumor, these profiling efforts have however not led to changes in treatment^2–4^. Chemotherapy, radiation and surgery have remained a standard of care for the past decades, and the overall survival for GBM patients stands at a dismal 16 months^2,5^. With the development of single-cell sequencing technologies it has become apparent that intratumor heterogeneity is widespread in GBM, as individual tumors consist of cells representing distinct subtypes or cellular states and contributing to therapeutic failure^6–8^.

Intratumor heterogeneity has been a hallmark of GBM even before modern sequencing technologies were available. The historical name “glioblastoma multiforme” referred to a non-uniform appearance of the cells with intermixed necroses and microvascular proliferation within a single histology section. The intratumor heterogeneity of GBM is also apparent in magnetic resonance imaging, which distinguishes these GBM from lower grade IDH1 mutant gliomas^9–12^. However, the degree to which analyses based on a single biopsy fail to capture this heterogeneity is not yet known.

Multi-region biopsy protocols have revealed the extent of clonal diversity across many cancer types, including GBM^13–15^. More recently, MRI-guided neuronavigation enabled the collection of multiple samples with their recorded position in the tumor, providing a first glance at the 3D spatial organization of GBM^16^. The capacity to obtain spatially distant tumor biopsies raises important questions about how these data should be used to deepen our understanding of the GBM ecosystem.

Another imaging modality showing GBM heterogeneity at the macro level is peri-operative staining with 5-aminolevulinic acid (5-ALA)^17,18^. This natural metabolite accumulates in GBM cells with defects in the heme biosynthesis pathway^19,20^. In GBM patients receiving an infusion of 5-ALA a few hours before the surgery, its metabolic product protoporphyrin IX accumulates in cancer cells and can be excited by blue light to produce fluorescence that helps the neurosurgeon more clearly delineate the tumor margins. While improving tumor resection precision, 5-ALA-guided surgery is not impermeable to GBM’s heterogeneity, with most tumors displaying regional variation in staining^18,21^.

Despite the widespread evidence for regional heterogeneity in GBM, it has not yet been extensively evaluated in the context of drug development. Typical drug screening campaigns test the compounds of interest on several cell lines, but established cell lines, such as U87-MG or U251, do not capture the intratumor heterogeneity of GBM and thus are a poor predictor of translation of such screening results^22^. Neurosphere cultures have been widely used in the 3D format, as these primary cultures maintain cellular plasticity and can be propagated *in vitro* as well as *in vivo*^22–24^. However, it is worth noting that the culture conditions might favor cellular phenotypes and select for particular features, thus losing the heterogeneity seen in patient tumors. With growing evidence of spatial cellular heterogeneity in GBM^16,25,26^, representing this aspect *in vitro* is likely to impact outcomes of drug discovery studies. Yet, it remains unclear whether neurospheres derived from different regions of the tumor would converge on a single phenotype due to lack of support from the tumor microenvironment and *in vitro* selection or if they would recapitulate the diversity of the biopsies from which they were derived from.

To address these challenges, we sampled multiple spatially distinct regions of 6 GBM cases, representing all 3 molecular subtypes of GBM^27,28^, and derived neurosphere cell lines from each region. Clonal analysis of these tumors confirmed their heterogeneous nature. Interestingly, the neurospheres derived from different areas of the same tumor retained their diversity with respect to 5-ALA metabolism, varying proliferative capacity, and distinct transcriptional programs also reflected in variable responses to cyclin-dependent kinase (CDK) inhibitors. We found that the cellular composition of the original tumor sample, predicted based on bulk transcriptomic data, can be linked to drug response outcomes derived from the respective neurosphere line. Thus, while GBM neurospheres represent a valid tumor model, their spatial origin and heterogeneity need to be accounted for to enable reconstruction of the GBM ecosystem in targeted therapeutic screening efforts.

## MATERIALS AND METHODS

### Human tissue samples

All experiments with use of human tumor tissue were approved by Scripps Research IRB protocol #IRB-18-7209 and Cleveland Clinic Martin Health IRB. Details of the patient cohort are provided in **Table S1**. Prior to the surgery, patients have been injected with 5-ALA, and all tumors used in the study were 5-ALA-positive based on fluorescence observed upon surgical resection. GBM diagnosis was confirmed by a board-certified neuropathologist on fresh frozen tissue at the Cleveland Clinic. Fresh GBM tumor tissue was collected directly from the operating room at the time of surgery. For each tumor sampling, MRI-guided navigation (Medtronic) was used to select the collection area, enabling us to spatially localize the biopsies, and location categories were assigned by the neurosurgeon. Tumor fragments were dissociated mechanically (with a piece being processed into formalin-fixed paraffin embedded block) and enzymatically with papain-based Brain Tumor Dissociation Kit P (Milteny Biotec, Cat#130-095-942). Red blood cells were removed by cell pellet incubation in RBC lysis buffer (155 mM NH_4_Cl, 12mM NaHCO_3_, 0.1 mM EDTA).

### Primary cell line generation and cell culture

GBM cells from dissociated tumor were seeded at 1 million cells per T25 flask with 5ml of Neurobasal media (Life Technologies, Cat#21103049), with 0.5% B27 Supplement (Life Technologies, Cat#17504044), 0.5% N2 Supplement (Life Technologies, Cat#17592048), 1% GlutaMAX Supplement (FisherScientific, Cat# 35-050-061), 20ng/ml EGF (Sigma, Cat# GF316), 20ng/ml FGF (Arcobiosystems, Cat#BFF-H4117) and 1% PenStrep (Fisher Scientific, Cat#15-140-122). Growth factors were added every 2 days. Accutase (Fisher Scientific, Cat#MT25058CI) was used to dissociate the spheres and passage the adherent cells.

U-87 MG cell line (ATCC, Cat#HTB-14) has been cultured in DMEM (Corning, Cat#10-013-CV) with 10% FBS (Avantor, Cat#89510-186) according to manufacturer’s protocol.

All cell lines were maintained in humidified incubators at 37°C, 4% CO_2_.

### Proliferation assay

The cells were seeded in 96 well plates at a density of 20,000 cells/well, in 6 replicates. Growth factors were supplied every 2 days and measurements were performed every 4 days starting at day 2. Cell Titer 96® AQueous One Solution (Promega, Cat# G3580) was added at 20μL/100μL total volume and incubated for 2-3 hours. Absorbance was measured at 490nm.

### Drug treatment and dose response modeling

The cells were seeded at a density of 80-90,000 cells/well in 96-well plates, in 4 replicates. After 24 hours incubation, the compounds were added (concentration range 0.1-10,000 nM; **Table S11**) and incubated for 3 days. Cell Titer 96® AQueous One Solution (Promega, Cat# G3580) was added at 20μL/100μL total volume and incubated for 2-3 hours. Absorbance was measured at 490nm.

Dose response curves were fit using the DRC R package v 3.0-1. For cases where a control was available, the drc function with a 4 parameter log logistic model was used with the following parameters: upperl = c(Inf,100, 100, Inf), lowerl = c(0.0001,0.0001,0.0001,0.0001), fct = LL2.4(names=c(’slope’,’min’, ‘max’, ‘ec50’). The area under the curve (AUC) was computed using the trapezoidal method with points in concentration range of 10-15 – 105, incrementing the exponent by 0.1. Because concentration is used, a higher AUC corresponds with increased resistance while a low AUC indicates sensitivity.

### 5-ALA-based cell sorting

Fresh human tumor cells were sorted on BD FACSAria Fusion, based on their endogenous 5-ALA metabolite fluorescence, as patients received 5-ALA infusion before tumor removal. In experiments where cultured cells were used, sorting was performed 12h after addition of 10uM 5-ALA (Sigma, Cat#A7793) to the culture media. Gating was set based on a positive control, U87 human glioblastoma cells (ATCC), incubated for 12h with 10 uM 5-ALA. Sorted cells were processed either through single cell or bulk sequencing protocol.

### Single cell RNA sequencing and data analysis

Library preparation for scRNA-Seq was performed according to 10x Genomics Chromium 3’ v2 protocol, targeting 2,000 cells per sample. Demultiplexing and alignment of data were performed using STAR aligner 2.7.9a and the downstream processing was performed with Seurat^29^. The reads were aligned to Homo sapiens GRCh38 release 104 reference genome. The output of the pipeline was imported into R 4.3.3 with Seurat package 5.0.3^29^. The data was subset to include cells with 200-2500 transcripts and <5% mitochondrial gene content and normalized using “LogNormalize” method.

### Bulk RNA sequencing and analysis

For bulk RNA-Seq, RNA was extracted from a snap-frozen tissue or cell line pellets with TRIzol Reagent (ThermoFisher, Cat#15596026),), ZR BashingBead Lysis Tubes (Zymo Research Cat#S6012-50) and Direct-zol RNA Mini-Prep Plus kit (Zymo Research, Cat#R2071). Libraries were prepared using NEBNext Ultra 2 kit with rRNA depletion module1 (New England Biolabs) and sequenced on Illumina NextSeq 2000 platform, according to standard protocol. The reads were processed using the nf-core/rnaseq v3.16.0 pipeline using Nextflow v24.04.4. The command executed was nextflow run nf-core/rnaseq -profile singularity -c {nextflow_config} --input {samplesheet.csv} --outdir {output directory} --gtf {Homo_sapiens.GRCh38.104.gtf} --fasta {Homo_sapiens.GRCh38.dna.primary_assembly.fa} --aligner star_salmon --save_reference --save_align_intermeds -r 3.16.0. Briefly, the nfcore RNA-seq pipeline includes adaptor trimming with Trim Galore!, read alignment with Star and quantification with Salmon and outputs raw reads as well as a Transcripts per Million (TPM) normalized reads. Comparative analyses were performed with GSEA^30^.

#### Cell type and cell state deconvolution

The CIBERSORT X^31^ online platform was used to estimate the cell type and cell state proportions of bulk RNA samples. It requires an external single cell or sorted bulk reference with defined cell types and a mixture file which contains info from the bulk RNA data that has been formatted to match the reference. Three different references were used. Sorted bulk cell types from Klemm et al.^32^ were used to identify CD45− cells, a single cell reference from Abdelfattah^33^ was used to identify detailed cell types, and another single cell reference from Neftel et al.^7^ to identify cell states. CIBERSORT X requires consistent normalization techniques between references and bulk mixture files. To accommodate this, the bulk mixture used with the Abdelfattah et al.^33^ reference was normalized by dividing counts by the total counts of the patient, multiplying by 10000, followed by adding 1 and taking the natural log. The bulk mixture used with the Neftel et al.^7^ reference was normalized using TPM. The bulk mixture for Klemm et al.^32^ was not normalized as the Klemm reference was also supplied as raw counts. In all three cases, the Ensembl transcript Ids of the bulk RNA data were converted to gene symbols using the org.Hs.eg.db R package. If the primary gene symbol mapping was not in the single cell reference, alternative aliases were applied. The single cell references files were then converted into CIBERSORT signature matrix files on the CIBERSORT X online platform with the following parameters: Disable quantile normalization: true, kappa: 999, q-value: 0.01, No. barcode genes: 300 to 500, Min. Expression: 1, Replicates: 5, Sampling: 0.5, Filter non-hematopoietic genes from signature matrix during construction: false.

In order to derive sample specific RNA expression of CD45− cells, the Klemm deconvolution was run using CIBERSORT X high-resolution mode using the Klemm signature matrix GEP file for batch correction in B-mode and with quantile normalization disabled. The other two mixture files were deconvoluted by the standard cell fraction imputation module using the appropriate signature matrix file with the following arguments: Batch correction: enabled, Batch correction mode: S-mode, Single cell reference matrix file used for S-mode batch correction: All_genes_200_per_celltype_SignatureMatrixFile.txt, Disable quantile normalization: true, Run mode (relative or absolute): absolute, Permutations: 500

#### Shannon’s equitability

TPM normalized counts from the nf-core pipeline were used to compute the Shannon’s equitability of each sample using the following formula:

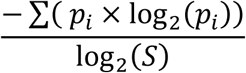

Where 𝑝_𝑖_ is the proportion of a single transcript within a sample and S is the number of unique transcript types in the sample.

### Exome sequencing and phylogenetic analysis

For whole-exome sequencing, DNA was extracted from a snap-frozen tissue with DNeasy Blood and Tissue kit (Qiagen, Cat# 69504) and shipped to Novogene for library construction with Agilent SureSelect Human All Exon V6 Kit and sequenced with Illumina NovaSeq 6000 platform, 200× coverage. Fastq files were then preprocessed using an implementation of the dna-seq-gatk-variant-calling pipeline (https://github.com/tjbencomo/ngs-pipeline, https://github.com/snakemake-workflows/dna-seq-gatk-variant-calling). The samples were aligned to the hg38 genome using bwa v0.7.17. Duplicates were marked using the MarkDuplicatesSpark command with GATK v4.2.0, followed by Base Quality Score Recalibration with GATK BaseRecalibrator and ApplyBQSR. Quality control metrics were performed by Samtools (v1.12) flagstat, fastqc v0.11.9, and mosdepth v0.3.1. A report was generated using MultiQC v1.11.

#### Variant calling

Variant calling was performed using Mutect2 v4.4.0 with --interval-padding 50, --genotype-germline-sites true, --genotype-pon-sites true --germline-resource set to af-only-gnomad.hg38.vcf.gz, --panel-of-normals {1000g_pon.hg38.vcf.gz}, and --intervals {SureSelect Human All Exon V6 r2 covered.bed file from the Agilent website}. GATK FilterMutectCalls with default parametes was run on the resulting vcf files to evaluate variant quality. Vcf files were then annotated with VCF2MAF (Cyriac Kandoth. mskcc/vcf2maf: vcf2maf v1.6. (2020). doi:10.5281/zenodo.593251) and VEP v104.3.

#### Constructing panel of normal for CNV calling

Following best practices for tumor-only CNV calling with PureCN v2.8.1, a panel of normal was constructed using processed matched blood WES samples from Mao et al^34^ which had been processed with Agilent SureSelect Human All Exon V6 Kit and NovaSeq 6000 platform. The 13 normal samples first underwent the identical preprocessing workflow as the tumor samples. PureCN coverage files were then constructed from the bam files by PureCN’s Coverage.R script, with default arguments. Germline variants were called using Mutect2 with identical parameters to the tumor variant calling but without using --panel-of-normals and --germline-resource, followed by FilterMutectCalls. The vcf files were merged with bcftools merge. A PureCN NormalDB file was created using PureCN’s NormalDB.R.

#### Calling CNV on tumor samples

Depth files were created for tumor samples using PureCN’s Coverage.R script, with default arguments. CNVs were called using PureCN.R using the coverage files and the panel of normals. An example command is below: Rscript /opt/PureCN/PureCN.R --out {output_directory} –tumor {coverage_loess_file} --sampleid {tumor_name} --vcf {mutect2_vcf_file} --stats-file {mutect2_stats_file} --fun-segmentation PSCBS --normaldb {normal_database} --mapping-bias-file {mapping_bias_file} --intervals {intervals_file} --genome hg38 --model beta --force --post-optimize --seed 123 --popaf-info-field POPAF.

#### Phylogenetic reconstruction

Due to the absence of matched normal tissue, a custom script was applied to combine the MAF annotation of the Mutect2 calling and the PureCN variant calling. Variants were removed if they did not have passed PureCN and Mutect2 variant calling filters, were hits in the germline database GnomAD_AF column at a greater than 1% allele frequency, or were silent or intron variants. PhylogicNDT v1.0 was run with the following arguments PhylogicNDT.py Cluster -i {patient} -sif {sif_file} --maf_input_type calc_ccf -rb --seed 4374 --maf --PoN 1000g_pon.hg38.vcf.gz –impute.

### Machine learning and predictive modeling

Sets of genes and pathways that can predict the response to dinaciclib in neurosphere lines were selected using the regression methods, Least Absolute Shrinkage and Selection Operator (LASSO) and Random Forest. LASSO was performed using the R package *glmnet* v4.1, by applying the maximum penalty to model coefficients; running 1,000-fold cross validations to find the optimal value of the tuning parameter, lambda that minimizes the mean squared error; and using the optimal lambda value to build a predictive linear model with the most important orthogonal genes or pathways. Random Forest-based feature selection was performed as previously described^35^, using the R packages *randomForest* v4.7 and *Boruta* v8.0. The gene sets identified by LASSO and Random Forest were used to build multiple linear regression models (MLR) using Prism v10.6.

To assess the impact of the predicted gene signatures on overall GBM patient survival, tumor sample and neurosphere-based MLR model scores were predicted for every patient in TCGA GBM dataset^36^. The predicted scores were then converted to z-scores, which were subsequently used to assign patients to low (below 25th percentile), moderate (between the 25th and 75th percentiles), or high (above the 75th percentile) scoring groups. Pairwise log-rank (Mantel-Cox) tests comparing the overall survival curves of these groups were performed using Prism v10.6.

## RESULTS

### Single-cell sequencing identifies 5-ALA^low^ tumor cell subpopulation

To test how spatially distinct GBM biopsies differ in their tumor and microenvironmental cell composition, we dissociated 6 biopsies from tumor OS1 into single cell suspension and leveraged the 5-ALA metabolite fluorescence to separate tumor cells (5-ALA^high^) from tumor microenvironmental cells (5-ALA^low^; **Figures 1A**, **1B and S1**). Sorted 5-ALA^high^ and 5-ALA^low^ populations were subject to bulk and single cell RNA sequencing, respectively. Single cell RNA sequencing (scRNAseq) identified distinct subpopulations of immune cells, including T/NK cells, macrophages/monocytes and dendritic cells (**Figures 1C**, **1D, S2A and S2B**). Within the 5-ALA^low^ population we also identified clusters of cells with transcriptional signatures of oligodendrocytes and astrocytes, as well as a population of cycling cells (**Figure 1D**). The frequency of different cell types in individual biopsies showed a gradient, with samples closer to the tumor surface enriched in macrophages/monocytes and dendritic cells, and samples located deeper in the tumor containing a larger proportion of astrocytes and oligodendrocytes (**Figures 1E and S2C**).

**Figure 1.**
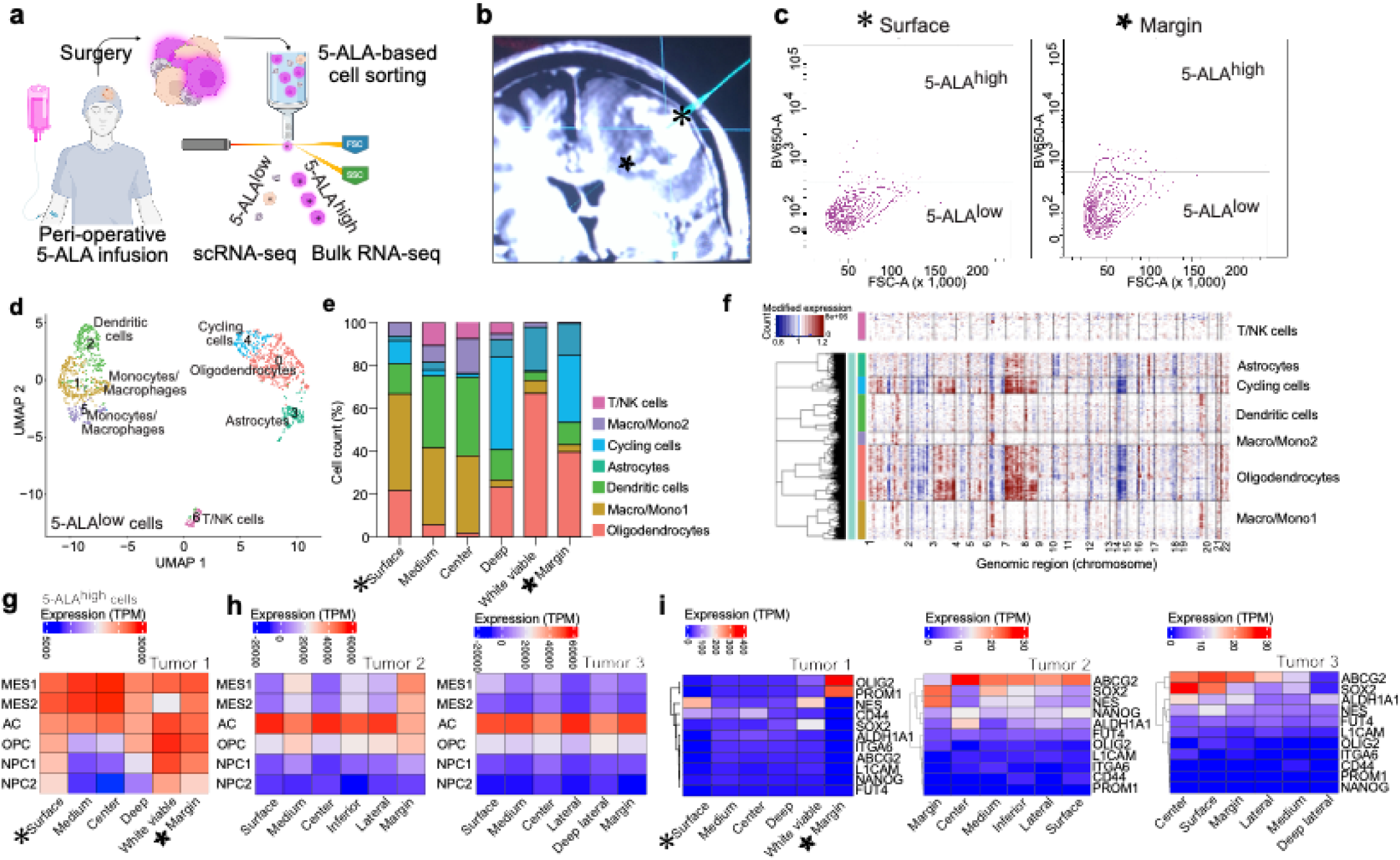
5-ALA-based sorting from multiregion biopsies reveals 5-ALA negative cancer cells enriched in deep tumor margins. **A**) Procedure outline. **B**) MRI neuronavigation image of GBM biopsies collected from Tumor 1. Stars denote surface and deep margin biopsies. **C**) 5-ALA-derived fluorescent cell sorting from surface and margin biopsies. **D**) Clustering of 5-ALA^low^ cells from 6 biopsies of Tumor 1 subject to singe cell transcriptomic profiling. **E**) Distribution of 5-ALA^low^ cell types across multiregion biopsies of Tumor 1. **F**) Copy number alterations across cell types identified in 5-ALA^low^ fractions of Tumor 1 biopsies inferred from scRNAseq. **G**) Expression of GBM cell state programs across 5-ALA^high^ fractions from multiregion biopsies of Tumor 1. MES – mesenchymal-like cells, AC – astrocyte-like cells, OPC – oligodendrocyte precursor-like cells, NPC – neural progenitor-like cells. **H**) GBM cell state signatures across 5-ALA high fractions of Tumor 2 and 3 (bulk RNA sequencing). **I**) Stem cell marker expression across multiregion biopsies from 3 tumors.

Since GBM cells can adopt transcriptional profiles of oligodendrocytes and astrocytes, we used our data to investigate whether the 5-ALA^low^ fraction could contain 5-ALA^low^ cancer cells. We applied computational algorithms to infer copy number alterations (CNA) from the scRNAseq transcriptomic data, with our T/NK cells as reference diploid cells. Indeed, we found that the clusters of cycling cells, as well as astrocytes and oligodendrocytes have multiple genomic regions with evidence of CNA based on InferCNV^6^ (**Figure 1F**) or CopyKat^37^ (**Figure S2D**). These results confirm that deep tumor margins contain subpopulations of metabolically distinct cancer cells, which may be missed during standard surgical resection. The divergence between distinct regions of the tumor was also apparent in our analysis of the 5-ALA^high^ fractions of the biopsies, with biopsies closer to the surface of the tumor displaying more mesenchymal-like cell state and higher levels of stemness markers (**Figures 1G-1I**). Therefore, development of *in vitro* models that capture this diversity is of high clinical significance. Since several cell lines of 5-ALA+ and 5-ALA− status have been previously developed^38,39^, we next sought to perform a comprehensive characterization of neurosphere lines derived from spatially distinct regions of GBM tumors, with varying 5-ALA-driven fluorescence status.

### A multi-region biopsy protocol enables derivation of neurospheres of distinct spatial origin

Our cohort for neurosphere derivation consisted of six GBM (IDH WT) cases, for which 5-ALA and MRI-guided neuronavigation was used to collect 4-8 biopsies per tumor from distinct tumor regions (total n=40 tumor biopsies; **Table S1**). The samples were classified as “central” or “peripheral” based on the MRI location. The samples most distant from the surgery entry point, buried deeper in the brain parenchyma, were annotated as “deep”. From each biopsy, neurosphere cultures were derived to assess whether spatial intratumor heterogeneity is reflected in these spatially divergent *in vitro* models.

First, we performed characterization of the original tumor samples. Based on transcriptomic profiling, tumors OS11, OS21 and OS29 represent proneural, mesenchymal and classical subtypes^27,28^, with all biopsies classified as the same subtype (**Figure 2A**). Tumors OS23 and OS27 were more heterogeneous with regards to this classification, clustering in between class centroids of the three GBM subtypes. Tumor OS28 was primarily classified as classical, with biopsy OS28-1, denoted as a subcortical tumor edge-containing normal brain, displaying a proneural transcriptional profile. Histopathological analysis showed that the biopsies had diverse levels of necrosis, thrombosis, immune infiltration, and cellularity (**Figure 2B**). When investigating whether these features could predict success in deriving fast-growing neurosphere cell line from the biopsies, we found no clear correlation between quantification of any of these features and the ability to derive neurosphere cultures (**Figure 2B, Table S2**). This finding suggests that even moderate to high levels of necrosis in a given biopsy can sustain a population of GBM cells able to grow in culture; moreover, a high mitotic index in the tumor tissue may not always be supported *in vitro*, as it may depend on cues from the tumor microenvironment, including interactions with neurons and glial cells^40,41^. Biopsies containing mostly normal brain cells and no features associated with the presence of the tumor mass gave rise to slow-growing neurospheres, suggesting that the rare GBM cells infiltrating the brain parenchyma may be intrinsically less proliferative than cells supported by the tumor microenvironment. Interestingly, computational deconvolution of cell types^31^ present across different areas of the tumor did not yield clear gradients between the tumor periphery and center (**Figure 2C**).

**Figure 2.**
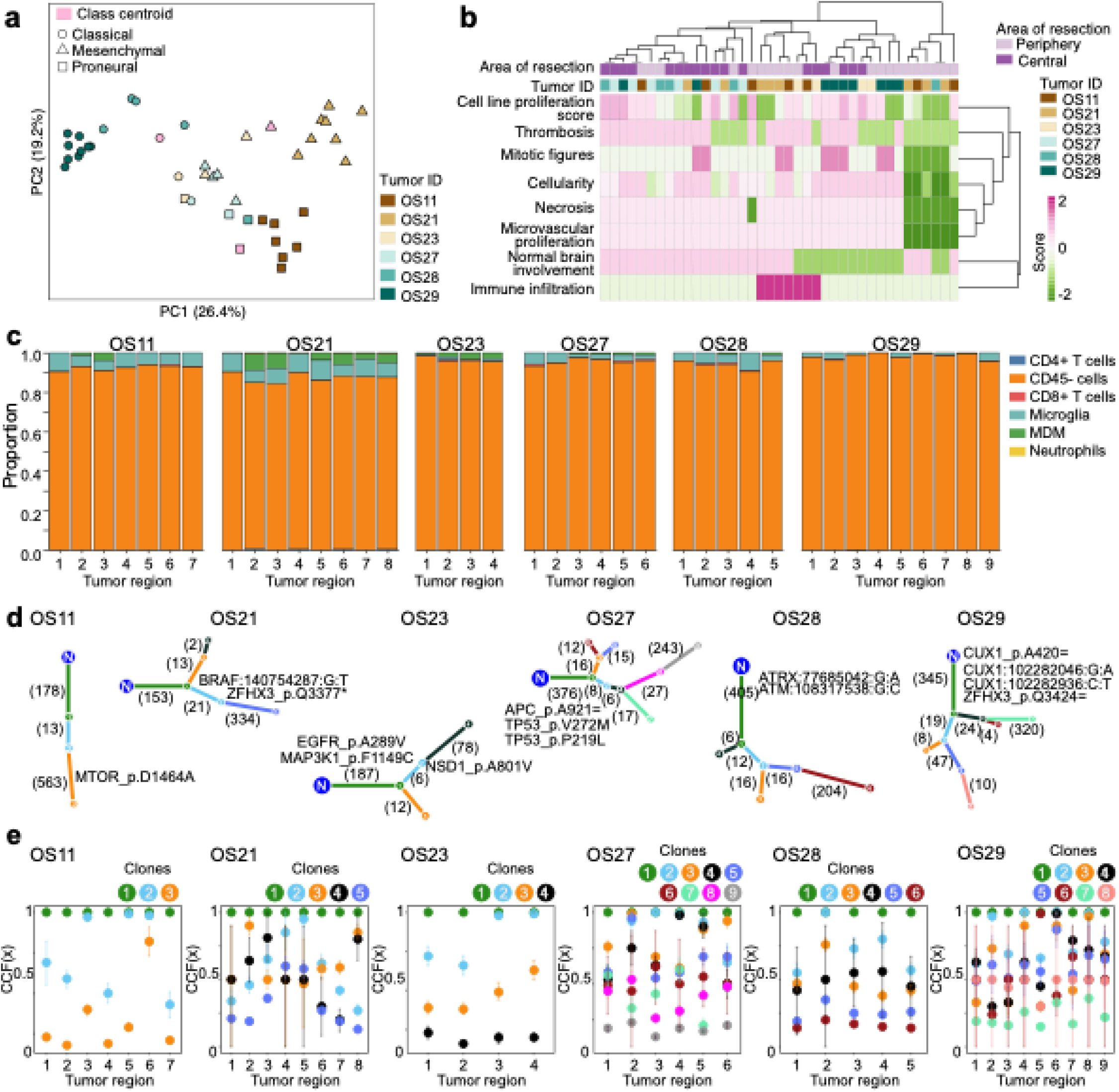
Intratumor heterogeneity across distinct regions of the GBM tumors. **A**) Subtype classification of multiregion biopsies. Subtype-specific signatures were used as class centroids (pink) and classification was based on principal component analysis. **B**) Histopathological analysis of multiregion biopsies based on H&E staining evaluation. **C**) Estimated frequency of cell types across multiregion biopsies based on CIBERTSORT X analysis of bulk RNAseq data using Klemm et al. cell-specific signatures. Numbers on x axis represent individual biopsies. MDM – Monocyte-derived macrophages. **D**) Phylogenetic deconvolution of clonal structure of 6 GBM tumors. Subclonal pathogenic mutations are denoted. **E**) Clonal composition estimation for multiregion biopsies. CCF – Cancer cell fraction. Error bars - confidence intervals for the frequencies. Numbers on X axis represent individual biopsies.

The acquisition of multi-region biopsies allows for inference of clonal composition of the tumors as well as their evolutionary trajectories^13,14,42^. We thus performed high coverage exome sequencing to reconstruct the clonal composition and ancestry of the tumors in our cohort. After correcting for tumor purity, we identified 3-9 clones present in each tumor (**Figures 2D**, **2E, S3 and S4, Table S3**). Three of the GBM cases were predicted to have more than one branching event. In two GBM cases, branching of a clone was associated with known pathogenic mutations (MTOR p.D1464A in OS11 tumor clone 3 and BRAF and ZFHX3 mutations in OS21 clone 5; **Figure 2D**). Interestingly, the frequency of the BRAF-driven clone 5 and its ancestral clone 2 in OS21 was significantly underrepresented in the tumor periphery (biopsies OS21-1 to 2 and OS21-6 to 8), at 0.13-0.27 clone 5 fractions, while it was highly abundant in the tumor center (biopsies OS21-4 and OS21-5; **Figure 2E**). In OS27, the phylogeny reconstruction predicted three branching events of clonal evolution, with clone 8 present at higher frequencies in the peripheral biopsies (OS27-1 and 6) and its progeny clone 9 present at low frequencies across all biopsies, which could be explained by increase migratory capacity of this new clone (**Figure 2E)**. Overall, these results indicate that clonal composition is variable across different locations in the tumor, consistent with prior whole genome studies demonstrating clonal divergence in GBM^15,16,43^.

### Neurospheres derived from spatially distinct tumor regions retain divergent phenotypes

Next, we focused on characterization of the neurospheres derived from spatially distinct regions of the tumor, to determine if they adopt a convergent phenotype in culture or if they retain the phenotypic diversity of the multi-region biopsies. From the 40 tumor biopsies described above, we derived a total of 30 primary cell lines, representing tumor margins, viable center, and deep margins (**Figure 3A**). Despite being subject to non-adherent conditions, our cultures displayed an array of mixed phenotypes, with some tumor regions producing round neurospheres and some having hybrid sphere and adherent populations (**Figure 3B**; throughout the manuscript we refer to all cultures as “neurospheres”). The extent of proliferation of the cell lines was also variable, with doubling times ranging from 4.8 to 18.6 days (**Figure 3C**), with OS21-7 being an outlier with minimal growth over 20 days after seeding in the proliferation assay. There were no apparent differences between the growth rates of the lines derived from the peripheral samples or the tumor center (**Figure S5**).

**Figure 3.**
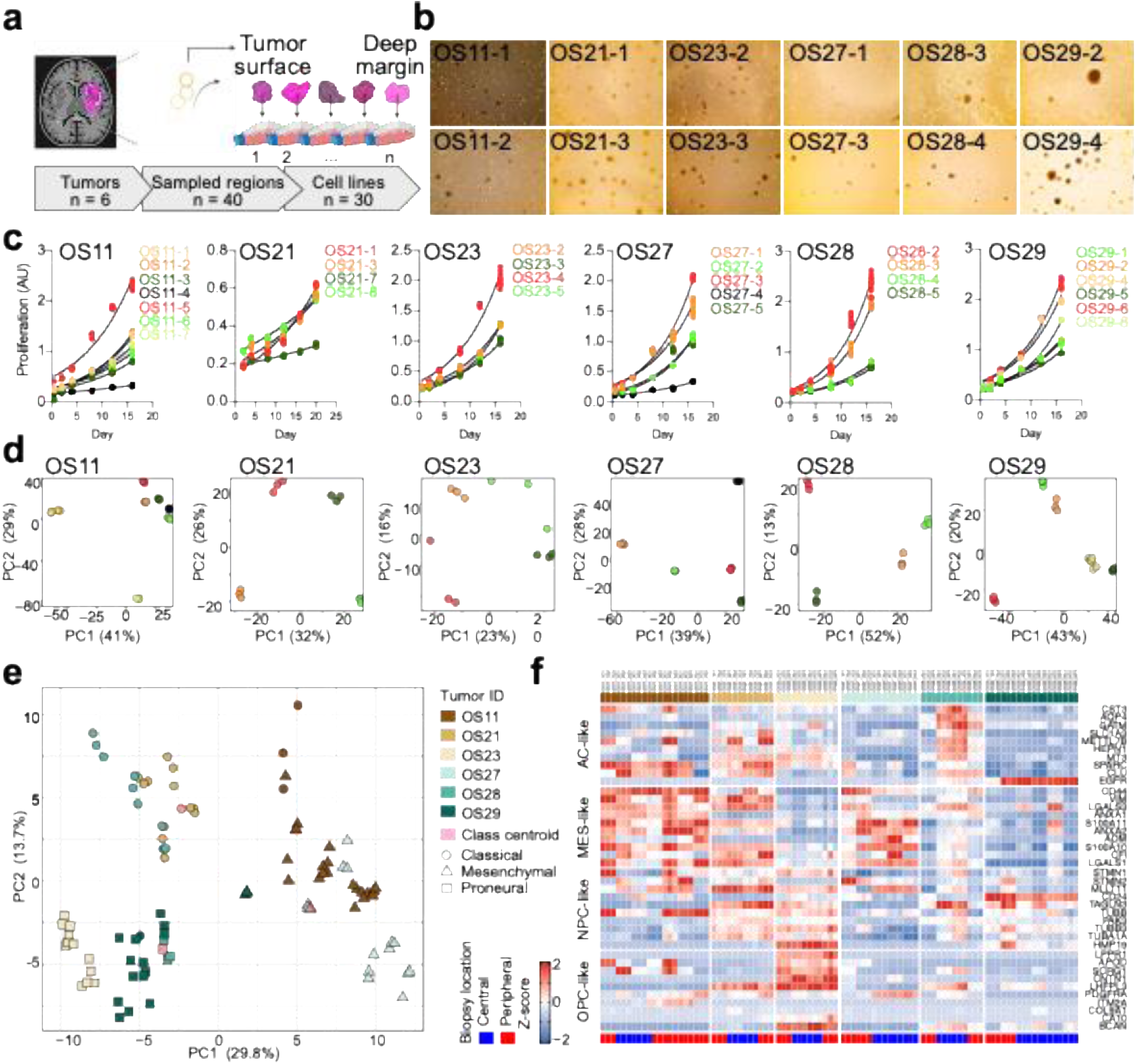
Neurosphere cell lines derived from multiregion biopsies remain heterogeneous in culture. **A**) Summary of the cell line derivation. **B**) Differential morphology of cells grown in non-adherent neurosphere-promoting conditions (passage 6-8). **C**) Proliferation of neurosphere cell lines derived from distinct regions of GBM tumors. **D**) Principal component analysis of neurosphere transcriptomes with different spatial origins. Color scheme corresponds to C). **E**) Subtype classification of multiregion biopsies. Subtype-specific signatures were used as class centroids (pink) and classification was based on principal component analysis. **F**) Cell state signature gene expression across the multiregion neurosphere cultures. MES – mesenchymal-like cells, AC – astrocyte-like cells, OPC – oligodendrocyte precursor-like cells, NPC – neural progenitor-like cells.

Transcriptional profiling of the patient-derived cell lines, performed in triplicates, showed that differences in proliferation were not the primary drivers of variation in gene expression between the lines derived from different regions of each tumor (**Figure 3D**). We found that the GBM subtype classification of the cell lines was more homogenous compared to the classification of the original biopsies (**Figure 3E**), likely due to the diversity of the tumor microenvironment involvement captured in the multi-region biopsy protocol. Of note, subtype classification is based on scoring subtype-specific gene signatures (**Table S4**). Assigning a single GBM subtype, based on the top scoring signature, seems to classify the neurosphere lines differently from their parental tumors of origin, but this classification is mostly driven by subtle changes in expression levels of the subtype-specific genes (**Figure S6A**). Nevertheless, in some cases (OS11, OS21 and OS29) these shifts seem to be directional, as the changes in subtype signature scores are similar across the lines derived from different regions of the same tumor. The expression patterns of the genes specific for four GBM cell states (neural-progenitor-like, oligodendrocyte-progenitor-like, astrocyte-like, mesenchymal-like) ^7^ displayed more divergence linked to the spatial origin of the neurospheres, compared to the GBM subtypes (**Figure 3F**). Cell lines derived from the peripheries of OS11 showed an enrichment of the neural progenitor-like (NPC-like) cell state, while those from periphery of OS28 show a depletion of the astrocyte-like (AC-like) state.

### Neurosphere culture transcriptional signatures remain divergent despite *in vitro* culture conditions

To investigate how well the neurospheres recapitulate the ITH of the spatially distinct biopsies, we performed analyses directly comparing the two sample sets. As expected, at the transcriptional level the neurosphere cultures cluster separately from their corresponding primary tumor biopsies, due to the fact that the latter contain cells of the tumor microenvironment and infiltrating normal brain (**Figure 4A and S6B**). Despite the variability in proliferation, the neurospheres derived from the same tumor are transcriptionally similar to each other and cluster according to their tumor of origin (**Figures 4A-4D**). To measure the transcriptional heterogeneity between the cell lines and their parental tumor biopsies, Shannon’s equitability scores were computed for each sample. Interestingly, using this measure, we found that the mesenchymal tumor samples were more heterogeneous than their corresponding neurosphere lines (**Figure 4E**). The mesenchymal GBM subtype has been linked with higher infiltration by immune cells^43^, and hence the lack of these cells in culture could explain more homogeneity of mesenchymal neurospheres. In contrast, the classical GBM subtype, characterized by EGFR overexpression/amplification and higher copy number variation, is rich in astrocyte-like cancer cells^27,43^, and cultures derived from this subtype were more transcriptionally diverse compared to their tumor samples of origin.

**Figure 4.**
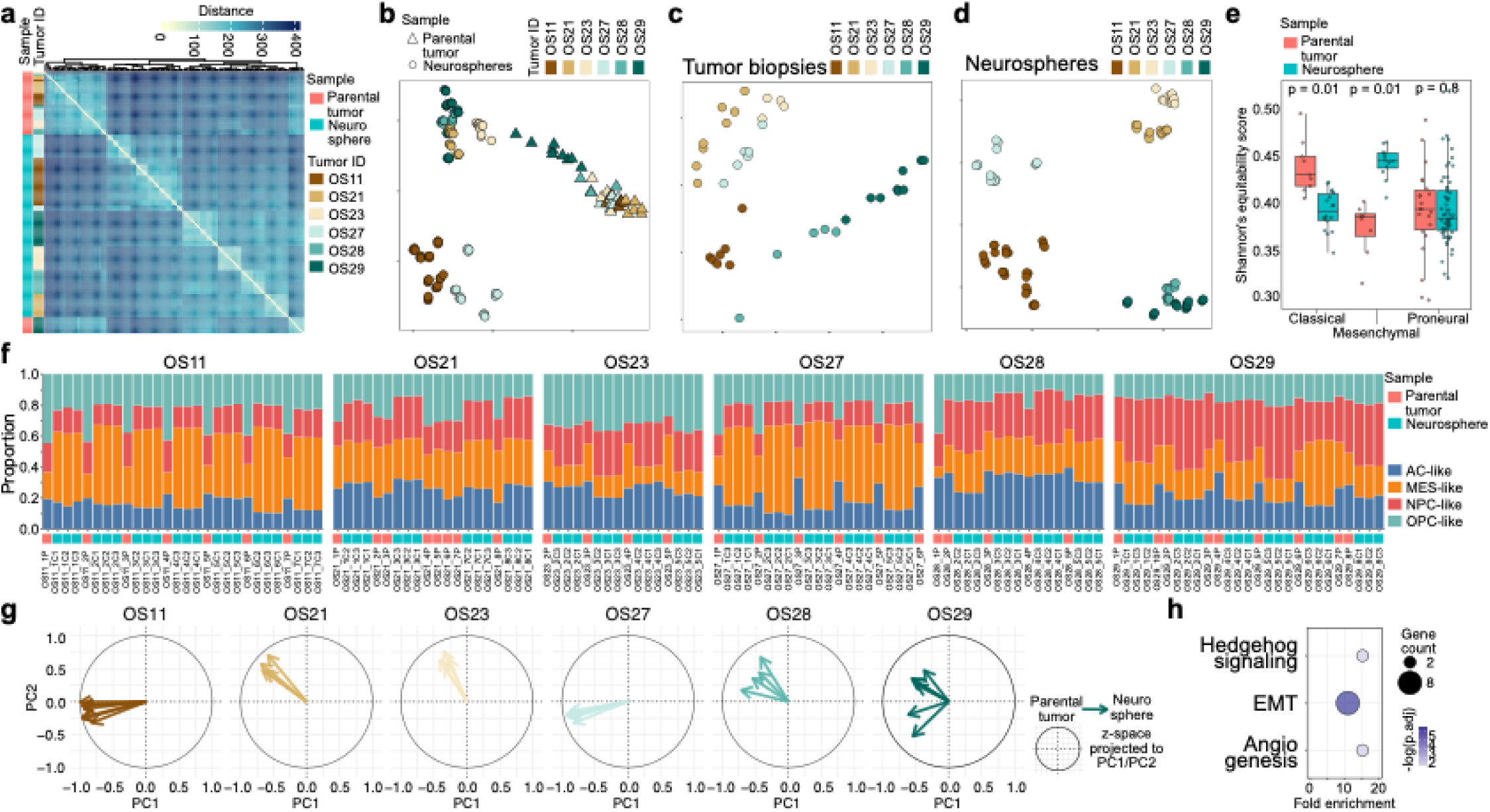
Transcriptional divergence of neurosphere cell lines from parental tumor samples. **A**) Correlation distance of transcriptional profiles of parental tumor biopsies and their corresponding neurospheres. **B**) Principal component analysis of the parental tumor biopsies and neurospheres. **C**) Principal component analysis of the parental tumor biopsies alone. **D**) Principal component analysis of the neurospheres alone. **E**) Shannon’s equitability score across GBM subtype-classified parental tumor biopsies and neurospheres. For neurospheres, their origin classification was used. P values of paired two-tailed t-test are shown. **F**) Estimated frequency of GBM cell states across multiregion biopsies and corresponding neurospheres based on CIBERTSORT X analysis of bulk RNAseq data. **G**) Parental tumor to neurosphere signature displacement (z-space projected to PC1/PC2). Arrow origin – parental tumor, arrow end – neurosphere. Angle of the arrow represents projection onto principal component 1 and 2. Radius represents the scaled magnitude of the z. **H**) Gene set enrichment in neurospheres compared to parental tumor samples after CIBERSORT-based signature correction (see Methods). EMT – epithelial-to-mesenchymal transition.

Previous studies reported that the mesenchymal phenotype is overrepresented in GBM cultures^44^. Consistent with this finding, a comparison of cell state signature distributions across distinct regions of the tumor and the corresponding neurospheres demonstrated that the mesenchymal phenotype expanded significantly in OS11 and OS27-derived lines, regardless of the spatial location (**Figure 4F**). However, in OS29 we observed an expansion of the NPC-like state at the expense of the AC-like state, while in OS21, the oligodendrocyte progenitor-like (OPC-like) state was depleted as the AC-like state expanded. Since GBM cell states are highly plastic and state transitions occur frequently, this transcriptional deconvolution may not clarify the relationships between the tumor and the corresponding cell lines. Therefore, we next set out to investigate how well the neurosphere lines represent the tumor regions they were derived from. To remove the contribution of the tumor microenvironment, which strongly differentiates the primary tumors from the cultured cells, we used computational deconvolution (see **Methods**). The tumor microenvironment-free (“TME-free”) signatures of the parental tumor were then compared to their corresponding neurosphere lines and projected as a transcriptional shift from the parental tumor to the neurosphere (**Figure 4G, Table S5**). The vectors representing the distinct regions of the tumor point in the same direction within the principal component space in 5 of our 6 cases, with OS29 neurospheres diverging in distinct directions from their primary origins. The parental tumor to neurosphere transcriptional signature shift does not seem to be the same across the tumors. These results suggest that the loss of the TME *in vitro* elicits distinct transcriptional adaptations for cells derived from different tumors, but similar for cells derived from distinct regions of the same tumor. The transcriptional shift from parental biopsy to neurosphere is supported by the overexpression of genes in the epithelial-to-mesenchymal transition (EMT), as well as NRP1, FSTL1 and SCG2, members of the Hedgehog and angiogenic pathways (**Figure 4H**). Thus, while mesenchymal phenotype enrichment is a common feature of GBM *in vitro* cultures, it may obscure the transcriptional features linked to the phenotypic and proliferative diversity of cells derived from distinct regions of the tumor and the underlying intrinsic properties specific to each tumor.

### Stemness, cell cycle and DNA repair pathways are upregulated in neurospheres derived from the tumor periphery

Increased cellular plasticity associated with EMT and expression of stemness markers allows cancer cells to quickly adapt to changes in their microenvironment and is thus responsible for drug resistance and tumor recurrence^45–47^. Therefore, we assessed the expression of stem-like state related genes across our cell lines. Based on our single-cell data (**Figure 1**), we hypothesized that the cell lines derived from the tumor periphery are likely to display a more stem-like phenotype. Indeed, in all cases except OS28, our neurospheres displayed a stemness gradient associated with the location from which they originated (**Figure 5A**). The neurospheres derived from samples localized deeper in the brain parenchyma in OS21, OS23 and OS27 expressed higher levels of stemness markers, while in OS11 and OS29 these markers were more abundant in lines derived from peripheral samples from the tumor surface. This finding suggests that the neurospheres derived from tumor peripheries retain higher levels of cellular plasticity and that these lines are likely more representative of the recurrence-driving populations in patients’ tumors.

**Figure 5.**
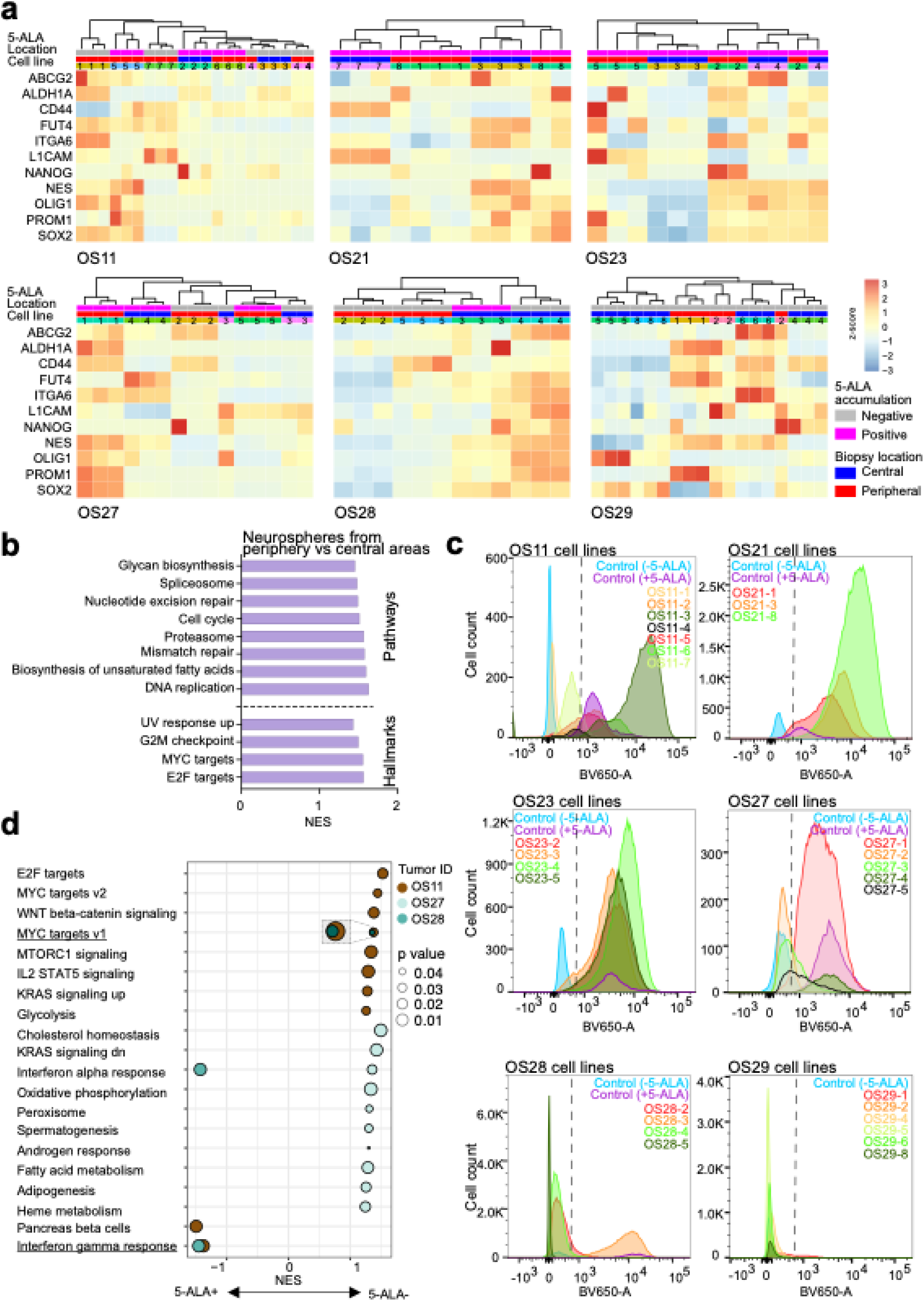
Gradients of stemness and location of origin contribute to differential gene expression and 5-ALA accumulation in multiregion neurospheres. **A**) Expression of stemness markers across neurosphere cell lines. **B**) Gene set enrichment analysis of genes highly expressed in neurospheres derived from the tumor periphery compared to those derived from central portions of the tumor. **C**) 5-ALA-derived fluorescence across neurosphere cultures. 5-ALA was added to the cultures in vitro. **D**) Gene set enrichment analysis of genes differentially expressed between patient-matched neurospheres with divergent 5-ALA accumulation status.

To learn more about the pathways driving the neurospheres derived from the tumor periphery and thus potentially driving local recurrence of the tumor, we next compared transcriptomic profiles of neurospheres derived from the central portion of the tumor with those derived from the superficial and deep margins. The neurospheres derived from tumor periphery had significantly upregulated cell cycle, DNA repair activity, biosynthesis of glycans and fatty acids signatures (**Figure 5B**). Thus, even though the neurospheres derived from the tumor center may divide faster, cultures representing tumor edges are highly active. Hallmark analysis revealed MYC targets and the G2M checkpoint as the primary drivers of these lines (**Figure 5B**).

### Tumor periphery-derived neurospheres retain differential ability to accumulate 5-ALA

Prior to surgery, all patients in our cohort received a 5-ALA infusion and the biopsies were collected from tumor regions displaying variable fluorescence of the 5-ALA metabolite. *In vitro* treatment of our neurospheres with 5-ALA also led to variable fluorescence, showing that metabolic heterogeneity of the distinct regions of the tumor is retained by our multi-region cultures (**Figure 5C**) and was not merely a reflection of the proliferation of the cultured cells (**Figure S5, Table S2**). Biopsies from tumors OS11, OS27 and OS28, representing all three subtypes of GBM, gave rise to sets of neurospheres with differential ability to accumulate 5-ALA. The extent of 5-ALA accumulation in GBM cells can be attributed to differential expression or activity of enzymes in the heme biosynthesis pathway, including FECH, HMOX1 and glutaminase 2, as well as transporters contributing to its flux, including ABCG2, ABCB6 and others^18,20,48^. Consistent with this observation, heme metabolism was significantly downregulated in 5-ALA^−^ lines derived from OS27 compared to the 5-ALA^+^ lines originating from the same tumor (**Figure 5D**). Pairwise comparison of 5-ALA^+^ and 5-ALA^−^ lines from the OS11 and OS28 tumors did not show convergence on the same pathways, except for MYC targets, which were upregulated in 5-ALA^−^ neurospheres from both tumors (**Figure 5D**). MYC overexpression has been linked to increased heme biosynthesis required for self-renewal and oncogenic transformation of hematopoietic progenitors^49^. Thus, it is likely that the differences in 5-ALA accumulation are driven by MYC. In contrast, 5-ALA^+^ lines from these two tumors displayed an enrichment of the interferon gamma response signature (**Figure 5D**), which can be linked to heme oxygenase-1 (HMOX1) expression and dampening of the anti-tumor immune response^50,51^.

Overall, our transcriptomic analyses of the neurosphere cells suggest that the lines representing the tumor periphery and those not accumulating 5-ALA are strongly associated with MYC activity.

### Drug response of neurospheres is linked to the tumor-of-origin’s phenotype

Our neurospheres derived from the tumor periphery were enriched in cell cycle and DNA repair gene signatures (**Figure 5B**). Since these cells are thought to drive local recurrence, targeting these tumor subpopulations could have a significant clinical benefit. To inhibit these pathways, we selected a panel of 7 drugs based on their recent evaluation in clinical trials for GBM patients and amenability for future medicinal chemistry studies and tested them in a 0.1nM – 10μM concentration range to assess their effect on neurosphere viability. The panel included the cyclin-dependent kinase (CDK) inhibitors dinaciclib and zotiraciclib^52^; the checkpoint kinase 1/2 inhibitor rabusertib, which induces cell cycle arrest^53^; and an inhibitor of the DNA damage repair pathway, AZD0156 (ATM inhibitor), as its brain-penetrant version has radiosensitizing effects in p53-mutant GBM^54^. We also tested ibrutinib, a Bruton’s tyrosine kinase inhibitor shown to impact the survival of cancer stem cells in GBM^55^; alpelisib, an inhibitor of PI3K, abnormally activated in most gliomas via mutations and dysregulation of receptor tyrosine kinases^56^; and rebastinib, which *in vivo* impacts angiogenesis by regulating ANGPT1/Tie2 expression, but can also inhibit proliferation and colony formation of GBM cells *in vitro*^57^. Temozolomide treatment was used as a reference, as this DNA alkylating chemotherapeutic agent was not expected to elicit potent effects when used as monotherapy, since its efficacy requires combination with radiation therapy. The response of the neurosphere cell lines to these treatments was variable between cells derived from distinct regions of the same tumor (**Figures 6A and S7, Table S6**). This variability could not be explained by differences in proliferation, as there was no significant correlation between doubling time and EC50 for any of the tested compounds (**Figure 6B**; Pearson’s correlation ranges from −0.3 to 0.2, with p values > 0.05).

**Figure 6.**
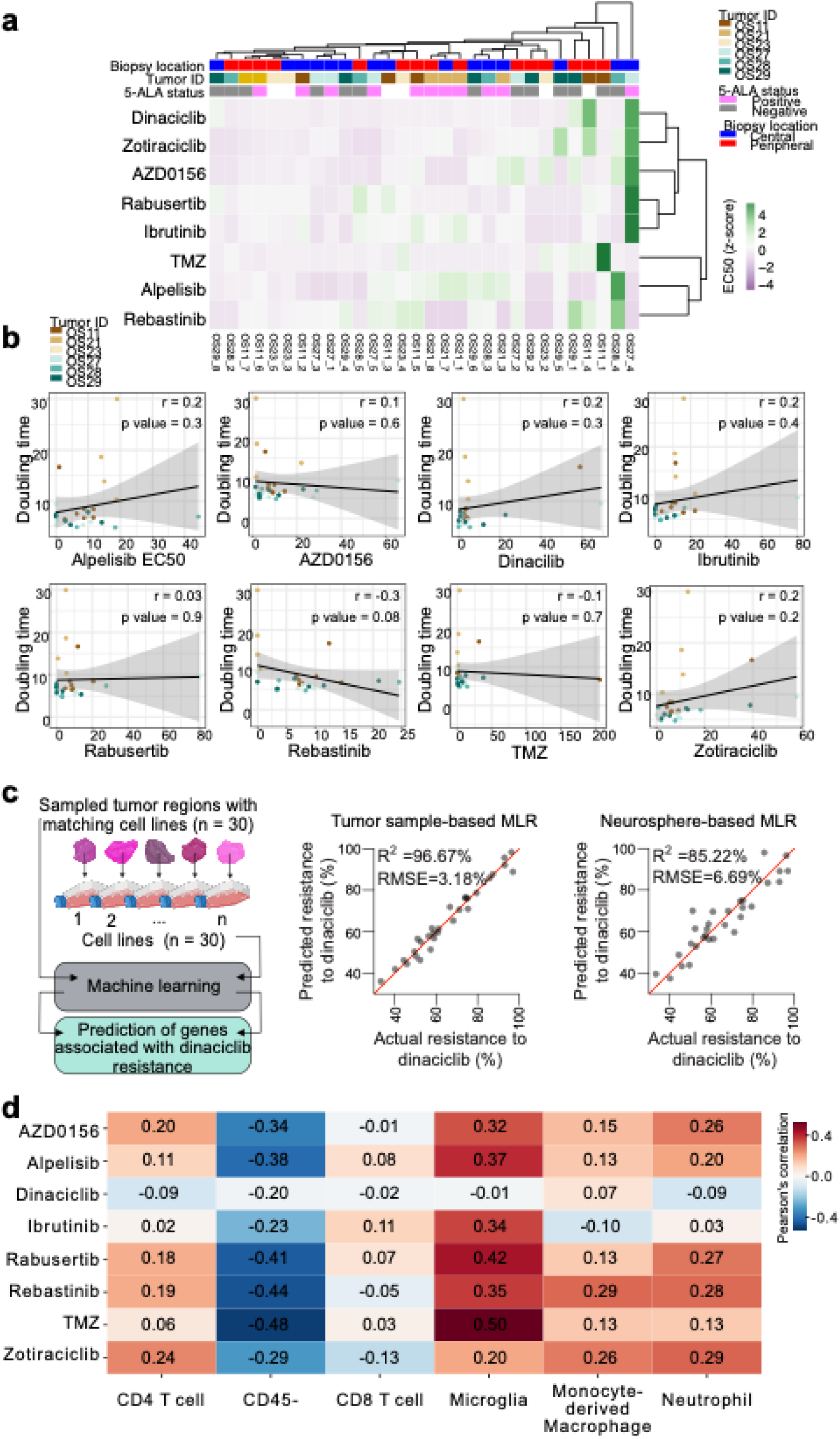
Differential response of cell lines to drug treatment linked to tumor phenotype. **A**) Drug sensitivity of multiregion neuropsheres. **B**) Correlation of drug sensitivity with neurosphere doubling time. Lines represent linear regression with shaded areas representing confidence intervals. None of the correlations reached a p-value < 0.05. **C**) Machine-learning-based prediction of dinaciclib response based on tumor sample or neurosphere transcriptomic data. MRL – multiple linear regression for combined LASSO and Random Forest predictions (see Methods). RMSE – root mean squared error. **D**) Pearson’s correlation of drug sensitivity (area under the curve, AUC) of the neurospheres with the predicted composition of the parental tumor biopsy. Low correlation values signify a lower AUC linked with higher frequency of a particular cell type in the original tumor biopsy.

To understand the basis for this variation, we turned to the machine learning approach. Given a relatively small dataset, we focused on dinaciclib response, as it was the most variable across our neurosphere lines. Least Absolute Shrinkage and Selection Operator (LASSO) regression and Random Forest-based feature selection were used to establish a prediction model for dinaciclib resistance based on either gene expression patterns across the tumor samples or across the neurospheres (**Figure 6C, Table S7-10**). The models identified 26 genes expressed in neurospheres as predictive of dinaciclib response (predicting 85.22% of the variance with root mean squared error of 6.69%), with RNA polymerase II subunit I (POLR2I) and SOX13 showing strong negative correlation with *in vitro* drug treatment outcome (**Figure 6C, Table S8**). The models built on expression profiles of tumor samples suggest that 23 genes, including inhibitor of CDK cyclin A1 interacting protein 1 (INCA1) and cell division cycle 23 (CDC23), when expressed in the tumor, correlate with *in vitro* dinaciclib response (**Table S9**). This model reached higher accuracy than the neurosphere expression-based model and predicted 96.67% of the variance in the response to dinaciclib, with a root mean squared error of 3.18% (**Figure 6C**). Interestingly, upregulation of pathways involved in neuronal differentiation in the original tumor samples was identified as a predictor of dinaciclib response *in vitro* (**Table S10**). Since the loss of CDK2 and CDK4 has previously been associated with a switch from proliferation to differentiation in neural stem cells^58^, it is possible that the tumors with a more neural profile are less sensitive to agents inducing this switch.

Inhibition of CDKs in GBM is currently being tested in clinical trials (i.e. NCT06810544, NCT06413706), and it will be interesting to test whether the signatures identified by our machine learning models could distinguish responders from non-responders in these trials. As a baseline, we noted that the low expression of the signature derived from the model trained on the tumor transcriptomic dataset and dinaciclib responses correlates with worse GBM patient survival (log-rank test p value = 0.0255, **Figure S8A-B**). The neurosphere expression-based model did not show a significant effect of the high vs low expression of the dinaciclib response-related genes in the untreated patient survival analysis (**Figure S8C**). It remains to be determined if the predictions based on the transcriptomes of tumor biopsies vs neurosphere cultures would perform differently in predicting CDK-targeting treatment response in patients.

Our results confirm that biopsies derived from distinct regions of the tumor may differ in their TME composition. It is plausible that the effects of cellular selection under distinct TME would result in distinct phenotypes of cancer cells. Although many of the transcriptional features driven by the TME might be lost in culture, we investigated whether our cell lines could still retain this TME-related specificity that could be mapped back to the tumor composition. To assess this question, we correlated the composition of the tumor region based on transcriptional deconvolution in **Figure 2C** with corresponding cell line drug response area under the curve (AUC). In this analysis, negative correlation signifies that a higher frequency of a particular cell population in the tumor biopsy was linked to a more effective *in vitro* killing of cancer cells derived from that tumor (**Figure 6D**). We found that a large frequency of glioma cells in the original tumor biopsy (CD45^−^ cells) was linked to better response of the corresponding cell line to temozolomide, while microglia abundance in the biopsy correlated with worse *in vitro* outcome. Interestingly, if the original tumor sample was enriched in pericytes, the corresponding cell line responded better to PI3K inhibition with alpelisib (**Figure S7B**). A large frequency of myeloid cells was weakly correlated with poor cell line response to all tested compounds. Similar analysis performed using GBM cell states instead of TME cell types did not render any significant correlations with cell line drug response. These results suggest that, at least in part, the transcriptional programs retained by GBM cells *in vitro* reflect their history of selection in distinct microenvironments in the tumor.

## DISCUSSION

The spatial organization of tumor tissue has recently emerged as an important aspect of heterogeneity within tumors^25,59–61^. On the meso scale, within a single tumor biopsy, hypoxic/necrotic foci appear to drive a layered organization of cellular states^26^. On the macro scale, across the whole tumor, analysis of spatially distant specimens revealed regional specificity in GBM and its microenvironment programs^16^. Our study provides the first glimpse into the tumor-wide diversity of phenotypes that can be propagated *in vitro*.

Despite stable culture conditions in non-adherent flasks and supplementation of epidermal growth factor (EGF) and fibroblast growth factor (FGF), the neurospheres derived from distinct areas of each GBM tumor in our cohort do not converge on the same phenotype and remain distinct from each other at the transcriptional level, but also in their proliferative and 5-ALA accumulation capacity. This is consistent with prior reports of patient-derived GBM organoids retaining intratumor heterogeneity *in vitro*^24,62,63^. Interestingly, bulk transcriptomic profiling does not provide a clear connection between the gene expression signatures of the derived cell lines with their corresponding tumor biopsy. This is largely driven by the elimination of the tumor microenvironment *in vitro* and the potential evolution of the cultures over time. Our data shows that subtype switching across cultures derived from the same tumor is typically directional, suggesting that this process is not random. Advances in single-cell transcriptomics allowing for sample fixation and multiplexing open new possibilities for more detailed profiling of GBM biopsies and subsequent cultures, which will be needed to further our understanding of the *in vitro* model of divergence and evolution. Such studies, albeit requiring a lot of resources, would improve our ability to faithfully model highly heterogeneous GBM tumors.

Interestingly, while our *in vitro* cultures do not readily match their original tumor biopsies’ phenotypes, several of the drug responses measured *in vitro* correlate with specific elements of the cellular composition of the original tumor. This indicates that some of the traits differentially expressed between neurospheres derived from distinct regions of the same tumor must be stable and are likely selected by the interactions with the TME in the original tumor. Given a similar subclonal composition of the original biopsies based on whole exome sequencing, these stable traits that remain different between cultures derived from different tumor areas might be linked to epigenetic stability. With tools like PATH^64^, single-cell analysis of our cultures could resolve the questions of heritability and plasticity of these features. Understanding these connections and limitations of the *in vitro* expansion will aid in the improved translation of GBM targeting drugs developed in pre-clinical models.

Multiple prior studies focused on the identification of vulnerabilities of GBM cells residing in the tumor margins by profiling the tumor biopsies^39,43,65–67^. The increased resolution of the MRI-guided multi-regional sampling allowed us to collect and develop *in vitro* models from multiple tumor margins. While stemness programs seem to be gradually increasing towards one of the margins across our samples, the cell cycle and DNA repair are specifically upregulated in 5-ALA^−^ tumor cells, which typically reside in the deep margins. This contrasts with prior studies, where 5-ALA− regions only gave rise to poorly growing cells forming a monolayer in serum-free media^38^. Interestingly, treatment of our 5-ALA− cultures with cell cycle inhibitors or DNA repair inhibitors did not show clear dependence on these pathways in the standard short-term assay. Because of the variable doubling time of the cultures, it is possible that drug screening capturing changes over extended timeframes and accounting for inherent growth dynamics of each culture would need to be developed. Since the ability to metabolize or extrude 5-ALA allows the GBM cells infiltrating tumor margins to evade detection during fluorescence-guided surgery, larger efforts to identify vulnerabilities of these populations hold the potential to prevent tumor recurrence.

Our proof-of-principle small drug set shows that predicting responses based on the most differentially expressed genes and pathways is challenging. It is intriguing that machine learning-based models are better at linking the *in vitro* drug response to gene expression in the original tumor samples rather than in the neurosphere lines on which the *in vitro* drug screening was performed. It is plausible that the abundance of growth factors and nutrients in culture media relaxes the pressure on transcriptomic output, making the phenotype “noisier”. Despite this hypothesis, inherent features of the tumor biopsy can be captured when drug pressure is applied to these high-abundance culture conditions. The neurosphere lines developed in our study will serve as a valuable resource for future large-scale screens aiming to combat tumor heterogeneity with respect to the spatial origins of in vitro GBM models and their 5-ALA accumulation status and phenotypic plasticity.

## ETHICS

All experiments with use of human tumor tissue were approved by Scripps Research IRB protocol #IRB-18-7209 and Cleveland Clinic Martin Health IRB.

## ACKNOWLEDGMENTS

We thank the members of the Janiszewska lab and the Michor lab for their critical reading of this manuscript and for useful discussion. We thank Marianna Franco for help with data analysis and Dr. Christoph Rader for his expertise in setting up the drug response experiments. We thank the staff at UF Scripps Genomics Core (RRID:SCR_017827), Flow Core (RRID:SCR_027278), and University of Florida ICBR NextGen Sequencing Core (RRID:SCR_019152) for their expert technical support. Illustrations were created with BioRender.

## AUTHORSHIP

R.S and M.J. conceived the study. R.S. performed the experiments. Computational analysis was performed by J.G. and R. S. A.R.T. helped with flow cytometry analysis. B.O. helped with cell line maintenance. D.H. helped with drug response set up. J.C.N. and K.W.N. built the machine learning predictive models. C.V. performed the histological analysis. O.S. collected the human surgical specimen and annotated clinical data. T.O.M. advised on data analysis. F.M. and M.J. supervised the study. M.J. wrote the manuscript, with input from all authors.

## CONFILCT OF INTERESTS

F.M. is a co-founder of and has equity in Harbinger Health, has equity in Zephyr AI, serves as a consultant for both companies and serves on the board of directors of Exscientia Plc. R.S has equity in Sethera Therapeutics and a Stealth Startup. F.M. & R. M. declare that none of these relationships are directly or indirectly related to the content of this manuscript. All other authors declare no competing financial interests.

## DATA AVAILABILITY

Further information and requests for resources should be directed to and will be fulfilled by the lead contact, Michalina Janiszewska.

De-identified human whole genome sequencing data have been deposited at dbGaP and accession number is listed in the key resources table. They are available upon request if access is granted. To request access, contact dbGaP (dbGaP: https://dbgap.ncbi.nlm.nih.gov). In addition, public summary-level phenotype has been deposited at dbGaP (dbGaP: phs004410) and are publicly available as of the date of publication. The accession numbers are also listed in the key resources table. Transcriptomic data for tumor biopsies and neurospheres are also available at dbGaP (accession number phs004410.v1.p1). Single cell transcriptomic data are available at GEO (accession number PRJNA1418147).

The code used in this study is available at Michor lab GitHub (https://github.com/Michorlab/Multi-region-GBM)

