## Supplemental Figures for "Multi-region biopsies and patient-derived neurosphere cultures reveal spatial divergence in glioblastoma"

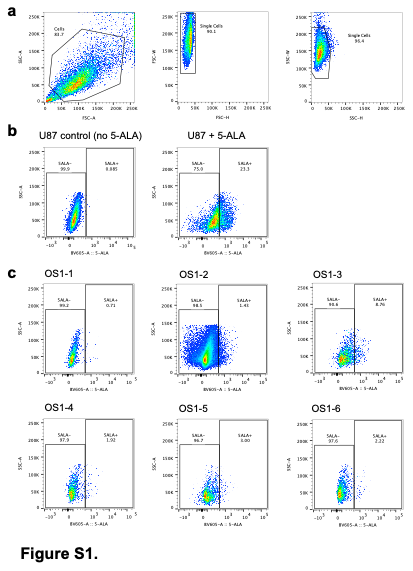

**Figure S1. Gating strategy for 5-ALA-based sorting. A**) Gating on live single cells. **B**) U87-MG cells were used as positive and negative controls for setting the gates for tumor cell sorting. Positive control was incubated with 5-ALA overnight. **C**) 5-ALA-based fluorescence gating for OS1 tumor sample sorting.

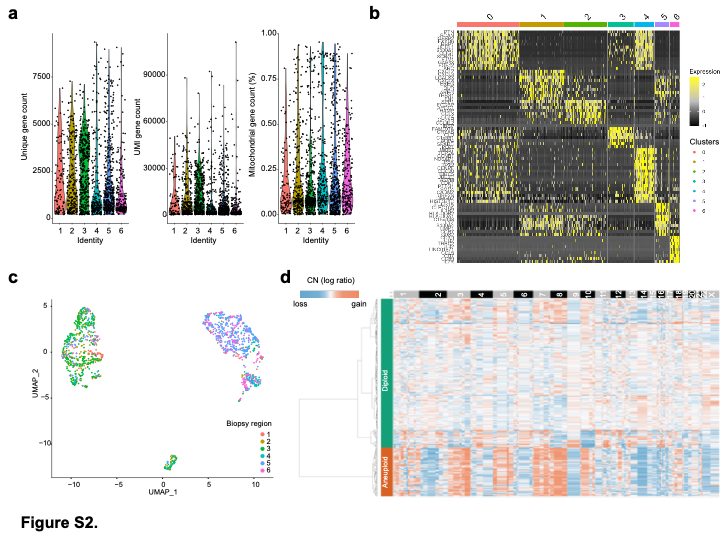

**Figure S2. Single-cell transcriptomic of 5-ALA- fractions QC. A**) Unique gene count, unique molecular identifier gene count and mitochondrial gene count per cluster identity**. B**) Genes differentially expressed between clusters. **C**) Cell clustering based on cell type colored by biopsy region of origin. **D**) Copy number and ploidy inference based on CopyKat algorithm.

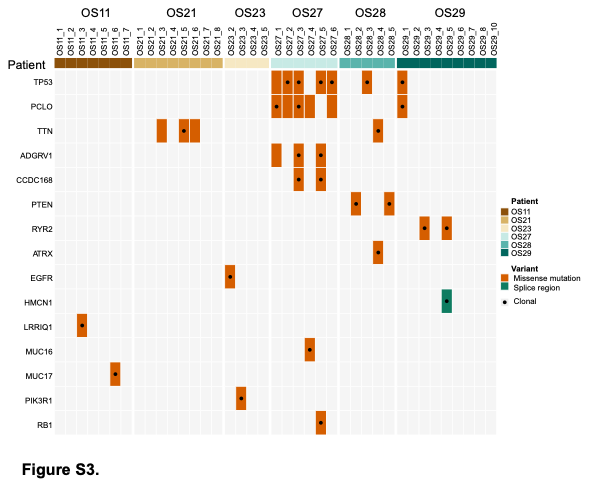

**Figure S3. Mutations in genes frequently affected in GBM across the multiregion biopsies.** Missense mutations and splice region mutations in genes frequently affected in GBM. Dot – clonal mutations. Only samples where at least one of the selected mutations was found are shown.

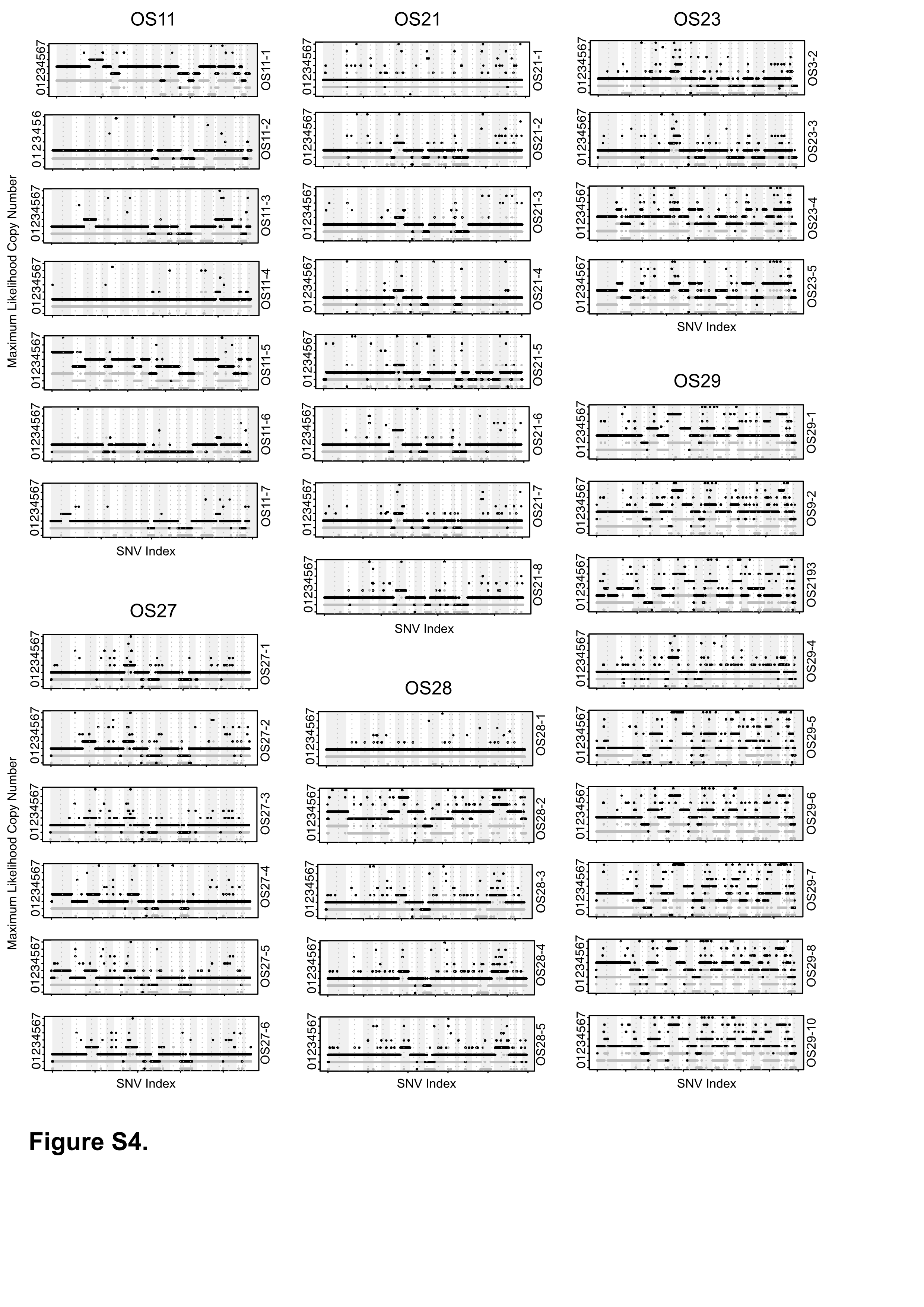

**Figure S4. Copy number variation across the multiregion biopsies.** Maximum likelihood copy number is plotted. SNV Index corresponds to chromosomal location (shaded areas indicate distinct chromosomes).

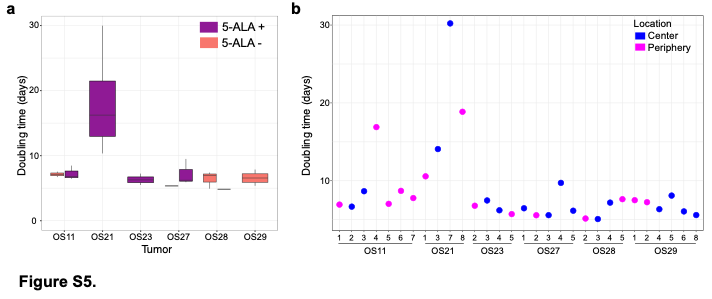

**Figure S5. Association of neurosphere proliferation with 5ALA status and location of origin. A**) Doubling time of all neurophere lines grouped by patient. Box plot represents quartiles, median and whiskers represent 5th and 95th percentile. **B**) Doubling time across all all neurosphere lines colored by the location of their biopsy of origin.

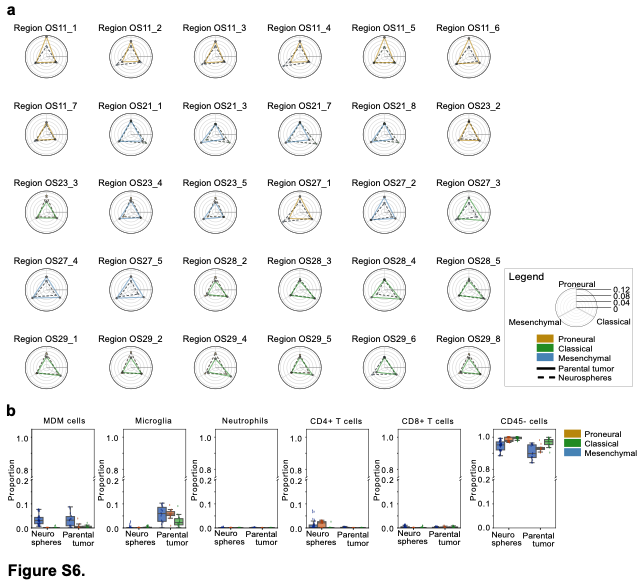

**Figure S6. Comparison of parental tumor biopsies and their corresponding neurosphere lines. A**) Subtype switching between parental tumor and cell lines. The tips of each triangle represent transcriptional score for mesenchymal, proneural and classical signature. Shaded area – parental tumor. Clear area, dotted lines – neurospheres. **B**) Estimated frequency of distinct cell types across neurospheres and parental tumors based on CIBERSORTx and Klemm et al. signatures. Box plot represents quartiles, median and whiskers represent 5^th^ and 95^th^ percentile.

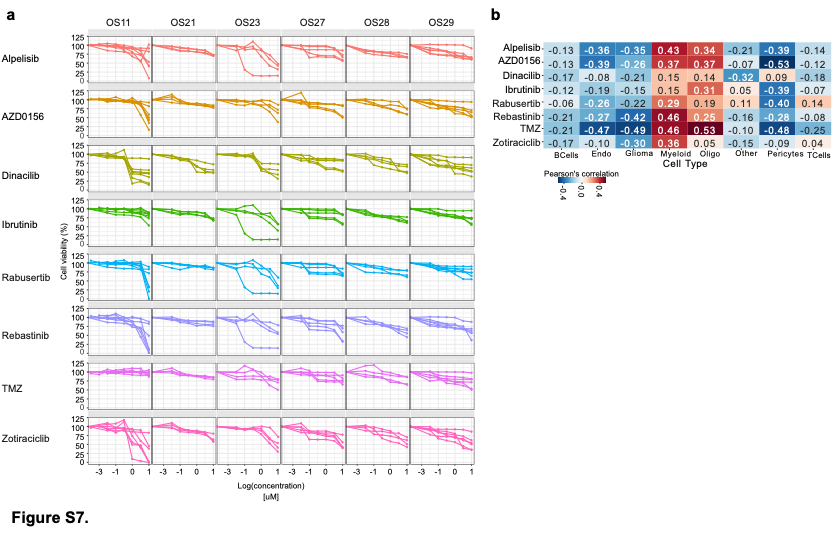

**Figure S7. Drug exposure in multiregion neurospheres. A**) Neurosphere drug responses, grouped by tumor. Cell viability was measured after exposing the neurospheres to the compounds. X axis represents concentration (log(X)) in uM. Normalized to DMSO. Each curve represents 5 replicates with standard deviation for each neurosphere line derived from a given tumor. **B**) Perason’s correlation of drug sensitivity (area under the curve, AUC) of the neuropsheres with the predicted composition of the parental tumor biopsy. Cell type-specific signatures from Abdelfattah et al. Low correlation values signify a lower AUC linked with higher frequency of a particular cell type in the original tumor biopsy.

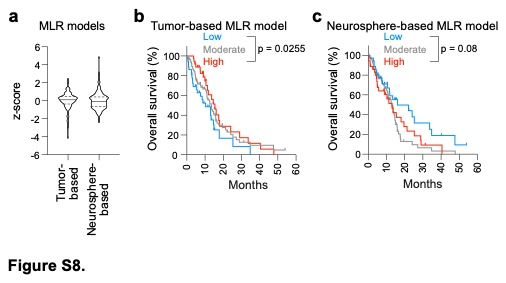

**Figure S8.** **Machine Learning-derived models across TCGA glioblastoma patients.** **A**) Distribution of multiple linear regression (MLR) model scores in TCGA glioblastoma patient samples (n=151, Brennan et al.)). Solid lines indicate the median, while dotted lines indicate the 25th and 75th percentiles. **B-C**) Overall survival of TCGA GBM patient cohort split by our tumors-based model scores (**B**) or neurosphere-based model scores (**C**). Low – <25^th^ percentile, moderate – 25^th^-75^th^ percentile, high - >75^th^ percentile. Log-rank test p values are shown.

**SUPPLEMENTAL TABLES**

**Table S1. Clinical data for the multiregion biopsy cohort.**

**Table S2. Cell line doubling time, 5-ALA status, and location of origin.** 5-ALA status was established by in vitro incubation with 5-ALA and fluorescence activated cell sorting; 1 - positive, 0 – negative.

**Table S3. Parental tumor exome sequencing-based purity and ploidy estimation.**

**Table S4. GBM subtype classification scores.** The scores for each GBM subtype were computed to assign the samples to a single subtype. Data related to Figure S5A.

**Table S5. Coordinates for transcriptional distance computed between parental tumors and cell lines.**

**Table S6. Drug response summary.**

**Table S7. Multiple linear regression of genes selected by LASSO or Random Forest to predict dinaciclib sensitivity.**

**Table S8. Predictive genes identified by machine learning from expression profiles in neurosphere cell lines.**

**Table S9. Predictive genes identified by machine learning from expression profiles of parental tumor samples.**

**Table S10. Predictive REACTOME pathways identified by machine learning.**

**Table S11. Compounds used in the drug sensitivity assays.**
